# Evaluation of different concentrations antibiotics on gut bacteria and growth of *Ectropis obliqua* Prout (Lepidoptera: Geometridae)

**DOI:** 10.1101/2021.04.14.439780

**Authors:** Tian Gao, Xiaomin Zhou, Rui Jiang, Chen Zhang, Song Liu, Tianyu Zhao, Xiayu Li, Yanhua Long, Yunqiu Yang

**Affiliations:** Anhui Agricultural University, State Key Laboratory of Tea Plant Biology and Utilization, Hefei, 230036, China; Anhui Agricultural University, School of Life Sciences, Hefei, 230036, China

## Abstract

The use of antibiotics to remove gut bacteria is a commonly used method to study gut function of insects. We assessed the impact of the artificial diet made from tea powder with different concentrations of antibiotics mixture (containing tetracycline, gentamicin, penicillin, and rifampicin) on *Ectropis obliqua* Prout (Lepidoptera: Geometridae) survival, growth, and reproduction. Antibiotic-induced bacterial clearance was monitored by Polymerase Chain Reaction (PCR) and by Colony-counting Methods. The results indicated that administration of the antibiotic mixture at a concentration of 200μg/ml caused the increase of gut bacteria, while a concentration of 300μg/ml was more effective in clearing gut bacteria but the concentration of 400μg/ml caused a large number of larval deaths. The concentration of 300μg/ml had no significant effect on the growth and development of *E. obliqua*, but had impact on fecundity. Therefore, we could use 300μg/ml concentrations of the antibiotic mixture to obtain sterile *E. obliqua* to study the function of intestinal microbes.

## Introduction

Insects are the most abundant class of animals worldwide, with more than one million species. More than half of insect species feed on plants; therefore, insects are the most important herbivores in the world[1]. Insect guts contain a substantial amount of microflora[2, 3] These gut microbes play an important role in mating and breeding; promoting digestion and growth, protecting insects from natural enemies, parasites, and pathogens; providing metabolic detoxification of toxins and insecticides; and enhancing host immunity[4-8]. For example, when *Drosophila melanogaster* flies are raised on only starch or only molasses as a food source, the adult flies only mate with members of the opposite sex that were raised on the same food source, But these preferences all disappeared after the antibiotic treatment[9]. By participating in the nitrogen cycle of the host to recycle the nitrogen from waste excreted, termite intestinal microorganisms can maintain the host’s nitrogen source balance[10]. Metagenomic analysis of the gut bacterial community of *Nasutitermes* revealed that the termites, which feed on plant xylem and phloem, host a large number of bacteria that can *Hamiltonella* defense, which colonizes aphids, directly kills the larvae of aphids’ natural enemies[11]. Other studies have addressed the relationship between insecticide resistance and gut microorganisms. For example, gut bacteria found in *Plutella xylostella* mediate resistance to chlorpyrifos[12]. *Citrobacter sp*. the gut symbiotic bacteria of the oriental fruit fly *Bactrocera dorsalis*, enhanced the host’s resistance to trichlorphon[13]. The bean bug *Riptortus pedestris* obtain *Burkholderia* that degrades fenitrothion from the soil, thereby obtaining resistance to the fenitrothion [14]. In short, gut microbial community play a significant role in the physiological activity of insects.

Previous studies have cleared gut bacteria using antibiotics to study the function of the gut microbiota. *Hypothenemus hampei* has difficulty degrading caffeine after being treated with antibiotics to remove its gut bacteria, therefore, and is unable to complete its life cycle [15]. When antibiotics were used to remove the indigenous midgut bacteria of the gypsy moth, *Bacillus thuringiensis* was unable to kill the gypsy moth larvae[16]. It is found that exposure to antibiotics significantly altered the honeybee gut microbial community structure and lead to decreased survivorship of honeybees[17]. *Rickettsiella* induce *Acyrthosiphon pisum* to form green pigments, which can prevent predation by natural enemies, and this ability disappears when *Acyrthosiphon pisum* are treated with antibiotics[18]. Some studies have explored whether antibiotics are toxic to insects. When *Spodoptera litura* were fed an artificial diet containing antibiotics to eliminate the gut microbiota, there was no adverse effect on growth, digestive enzyme activity increased, and detoxifying enzyme activity decreased[19]. Treatment with five antibiotics including tetracycline significantly reduced larval of Plutella xylostella growth and development, and eventually increased larval mortality and malformation of the prepupae[20]. Thus, there is a certain amount of controversy regarding whether or not antibiotics are toxic to insects. Therefore, the toxicity of antibiotic to the insect should be assessed prior to research the interaction between insect gut microbes and the host using antibiotic treatment. *Ectropis obliqua* Prout (Lepidoptera: Geometridae) is a major tea plant pest in Southeast China. Previous studies had focused on the biological characteristics of *E. obliqua*, the immunological effects of some of its proteins, and phylogenetic analysis of its mitochondrial genes[21, 22]. However, there had been few studies of E. obliqua gut bacteria composition and function. Obtaining sterile insects is based on studying the interaction between gut microbes and their hosts. The aim of this study was to screen out an appropriate antibiotic mixture concentration that could remove the gut bacteria of the *E. obliqua*, obtaining sterile insects, and not affect the growth and development of larval.

## Materials and Methods

### *E.obliqua* collection and rearing

*E.obliqua* moths were collected in June 2018 from a tea garden in Dongzhi (30°10′N, 117°02′E), Anhui province in Southeast China. The moths were rared on tea leaves in a climate-controlled insectary (23±2°C, 70%-80% relative humidity, and a photoperiod of 16:8 light:dark). The experimental larvaes were fed tea powder (TP) diet composed of 10g tea powder and 0.5g agar powder in 20ml deionized water. To prepare the TP diet, the agar powder and deionized water were mixed and heated to dissolve the agar powder, then cooled down to about 30 °C. Finally, adding tea powder made from fresh tea leaves (Baihaozao, tea variety), the artificial diet was completed. Previous reports indicated that treatment with a single antibiotic did not completely eliminate insect gut bacteria [20]. Therefore, we added different concentrations (200μg/ml, 300μg/ml, and 400ug/ml) of the mixture of four antibiotics including tetracycline, gentamicin, penicillin, and rifampicin to the artificial diet (TPA2, TPA3, TPA4). The manufactured artificial diet were then covered with plastic wrap and stored at 4 °C. All of the diet were used to rear *E. obliqua*.

### Assessment of *E.obliqua* growth and development

The first-instar larval of the *E. obliqua* were randomly selected and divided into 5 groups. There was five replications in each group and 10 individuals in each replication. The larvae was fed on 90mm petri dishes, each with ten larvaes. Artificial diet of E. obliqua supplemented with three concentrations of antibiotic mixtures, 200 μg/ml, 300 μg/ml, 400 μg/ml. Tea powder diet without antibiotic mixtures and the fresh tea leaves (TL) served as control. The experiments were carried at 23±2°C temperature and 70%-80% relative humidity along with photoperiod of 16:8 L:D. The number of dead larvae was recorded every day until all larvae were pupated. Observations were made daily on various biological indicators of *E. obliqua* viz. larval mortality, larval, pupal, egg, and adult development time, female and male pupal weight, pupation rate, fecundity, hatching rate, and eclosion rate.

### Cultivation of *E.obliqua* gut bacteria

To collect the gut contents, 20 4th-instar larvaes were randomly selected from the five groups, regardless of sex. The larvae were surface-sterilized with 75% ethanol for 90s and then rinsed with sterile deionized water. After dissection, the midgut contents were homogenized with 1 ml sterile deionized water and frozen at -80 °C. The stock solution of the larval gut homogenates was diluted 10,000 times, then 50 ml was used to inoculate liquid LB media (LB medium: NaCl 10.0 g/L, yeast extract 5.0 g/L, peptone 10.0 g/L, 2% agar powder);the step was replicated five times. The cultured gut bacteria were photographed after 48 hours of shaking at 150 rpm at 37 °C.DNA extraction and PCR amplification of the 16s rRNA V4 region.

To isolate DNA from the gut microbes of *E. obliqua* larvae, fifth-instar larvae (V-5, n=3) were surface-sterilized by dipping them in 75% ethanol once (for approximately 15s) and then rinsing them twice in sterile water rinse (for approximately 30s). Dissecting scissors were used to make a lateral cut behind the head capsule, and the gut was removed from the cuticle with larval forceps. The entire gut, including gut contents, was collected and placed in a 2.0-ml micro-centrifuge tube for processing (all steps were performed on ice). Samples were placed in -80 °C freezer. Total genomic DNA was extracted from the samples using a QIAamp DNA Stool Mini Kit (Qiagen, USA), using 1% agarose gel electrophoresis to detect the concentration and purity of DNA. The DNA was diluted to 1 ng/μL in sterile water. A 450-bp fragment from the 16s rRNA V4 region was amplified using the specific primer pair 27F-1492R (5’-AGAGTTTGATCCTGGCTCAG-3’, 5’-GGTTACCTTGTT ACGACTT-3’). All PCR reactions were carried out with Phusion High-Fidelity PCR Master Mix (New England Biolabs). The PCR products were mixed with an equal volume of 1× loading buffer (containing SYB green) and electrophoresed on a 2% agarose gel for detection. Samples showing a bright band of 400-450 bp were chosen for further experiments. The PCR products were diluted to equivalent concentrations and purified with a Qiagen Gel Extraction Kit (Qiagen, Germany).

### Quantitative PCR

Total RNA from two groups of samples were extracted using an SV total RNA isolation system with a DNase purification step (Promega) according to the manufacturer’s instructions. The 2-μg RNA sample was reverse transcribed using the PrimeScript™ RT Master Mix (Takara, Shiga, Japan). Quantitative real-time PCR (qPCR) was performed with using an ABI 7300 Real-Time PCR System (Applied Biosystems, Foster City, CA, USA) and GoTaq qPCR Master Mix (Promega) in a volume of 10 μL. The PCR conditions were as follows: 95 °C for 30 s; 95 °C for 5 s and 60 °C for 30 s, 40 cycles. The qPCR data was collected and analyzed via the 2^−ΔΔCt^ method[23]. All samples were independently measured three times. The forward and reverse primers used for the genes of interest have been reported previously. Gene expression was normalized to the CYP4G55v3 gene.

### Statistical analysis

To compare differences in means, one-way analysis of variance, with Tukey’s test at P ≤ 0.05 was performed. SPSS software for windows version 22.0 (IBM Corp Version 22.0, IBM SPSS Statistics for Windows; IBM, Armonk, NY, USA) and Prism 5 (GraphPad Software, La Jolla, CA, USA) software were used to perform the statistical analysis.

## Results

### Effects of different concentrations of antibiotic diets on *E.obliqua* gut bacteria and survival

The first-instar larval of *E.obliqua* was fed on different treatments’ diets (TL, TP, TPA2, TPA3, TPA4) until pupation. During this period, the larval of mortality and various physiological indicators were recorded every day. Compared with other treatment diets, feeding the 400μg/ml(TPA4) of antibiotic mixture diet on larvals of *E.obliqua*, the survival rate of the larval was significantly reduced. However, compared with TL and TP groups, feeding 200μg/ml(TPA2) or 300μg/ml(TPA3) of antibiotic mixture diet on larvals, the survival rate of the larval was no effect (Fig 1). PCR-Agarose gel electrophoresis analysis confirmed that treatment with 300μg/ml or 400μg/ml of the antibiotic mixture more lowly gene expressed and significantly reduced the number of gut bacteria in *E.obliqua* larvae (Fig 2). However, treatment with 200μg/ml of the antibiotic mixture resulted high gut microorganisms gene expression. CFU analysis revealed that the number of cultivable gut bacteria significantly reduced, as increasing the concentration of antibiotic mixture. When using 300μg/ml of the (Table 1)antibiotic mixture diet treated, the gut bacteria was effectively eliminated by antibiotics.

**Table 1.**
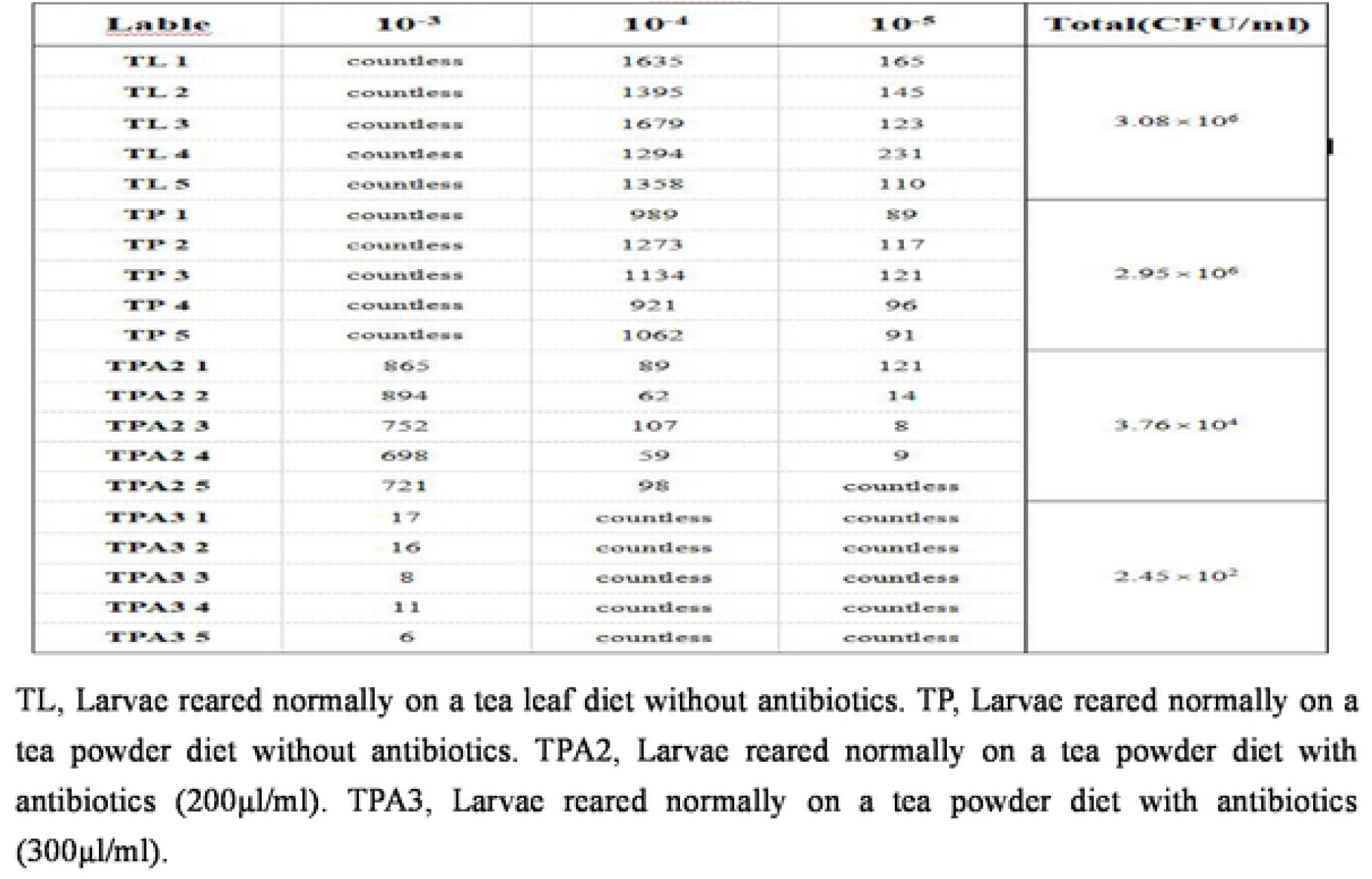
Culture of gut microorganisms in *E.obliqua*.

| Lable | 10<sup>-3</sup> | 10<sup>-4</sup> | 10<sup>-5</sup> | Total(CFU/ml) |
| --- | --- | --- | --- | --- |
| TL 1 | countless | 1635 | 165 | 3.08 × 10<sup>6</sup> |
| TL 2 | countless | 1395 | 145 |  |
| TL 3 | countless | 1679 | 123 |  |
| TL 4 | countless | 1294 | 231 |  |
| TL 5 | countless | 1358 | 110 |  |
| TP 1 | countless | 989 | 89 | 2.95 × 10<sup>6</sup> |
| TP 2 | countless | 1273 | 117 |  |
| TP 3 | countless | 1134 | 121 |  |
| TP 4 | countless | 921 | 96 |  |
| TP 5 | countless | 1062 | 91 |  |
| TPA2 1 | 865 | 89 | 121 | 3.76 × 10<sup>4</sup> |
| TPA2 2 | 894 | 62 | 14 |  |
| TPA2 3 | 752 | 107 | 8 |  |
| TPA2 4 | 698 | 59 | 9 |  |
| TPA2 5 | 721 | 98 | countless |  |
| TPA3 1 | 17 | countless | countless | 2.45 × 10<sup>2</sup> |
| TPA3 2 | 16 | countless | countless |  |
| TPA3 3 | 8 | countless | countless |  |
| TPA3 4 | 11 | countless | countless |  |
| TPA3 5 | 6 | countless | countless |  |

TL, Larvae reared normally on a tea leaf diet without antibiotics. TP, Larvae reared normally on a tea powder diet without antibiotics. TPA2, Larvae reared normally on a tea powder diet with antibiotics (200µl/ml). TPA3, Larvae reared normally on a tea powder diet with antibiotics (300µl/ml).

**Figure 1.**
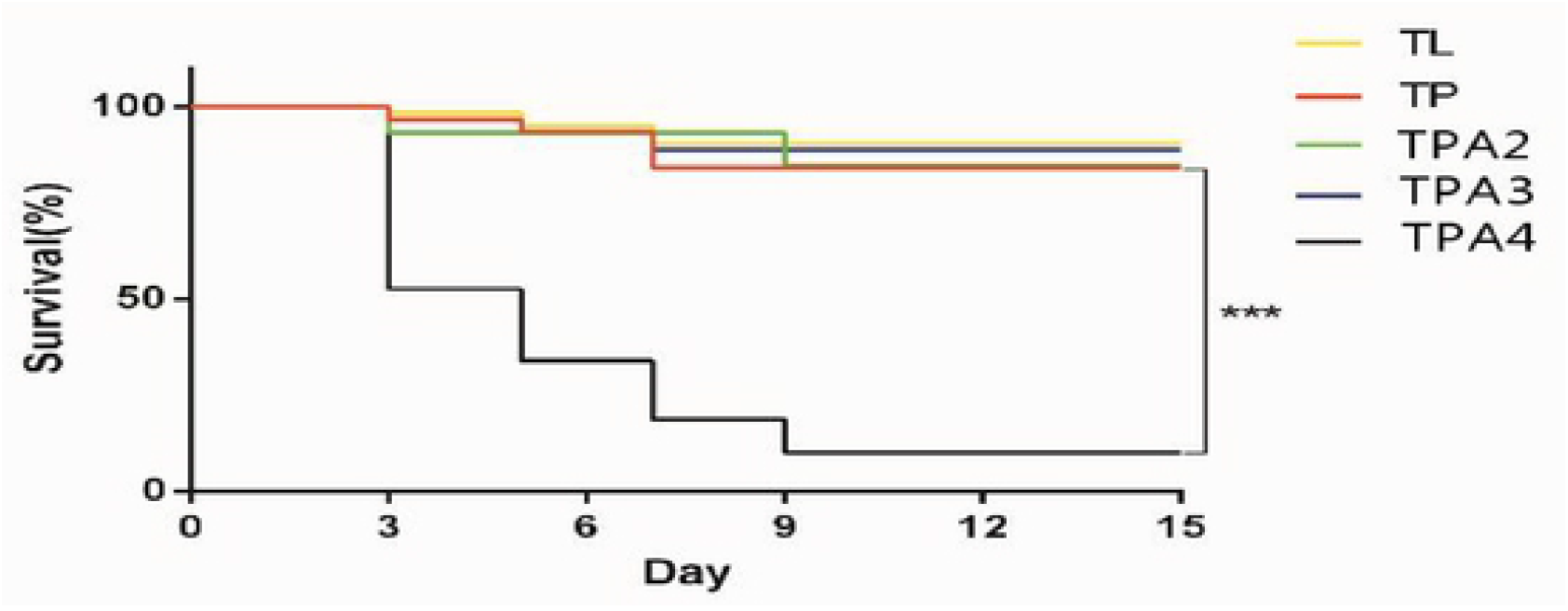
Survival rates of *E.obliqua* after feeding with different diets. TL, Larvae reared normally on a tea leaf diet without antibiotics. TP, Larvae reared normally on a tea powder diet without antibiotics. TPA2, Larvae reared normally on a tea powder diet with antibiotics (200µ1/ml). TPA3, Larvae reared normally on a tea powder diet with antibiotics (300µ1/ml). TPA4, Larvae reared normally on a tea powder diet with antibiotics (400µ1/ml). Using t-test to assess differences between groups, *p<0.05;**p<0.01;***p<0.0001.

**Figure 2.**
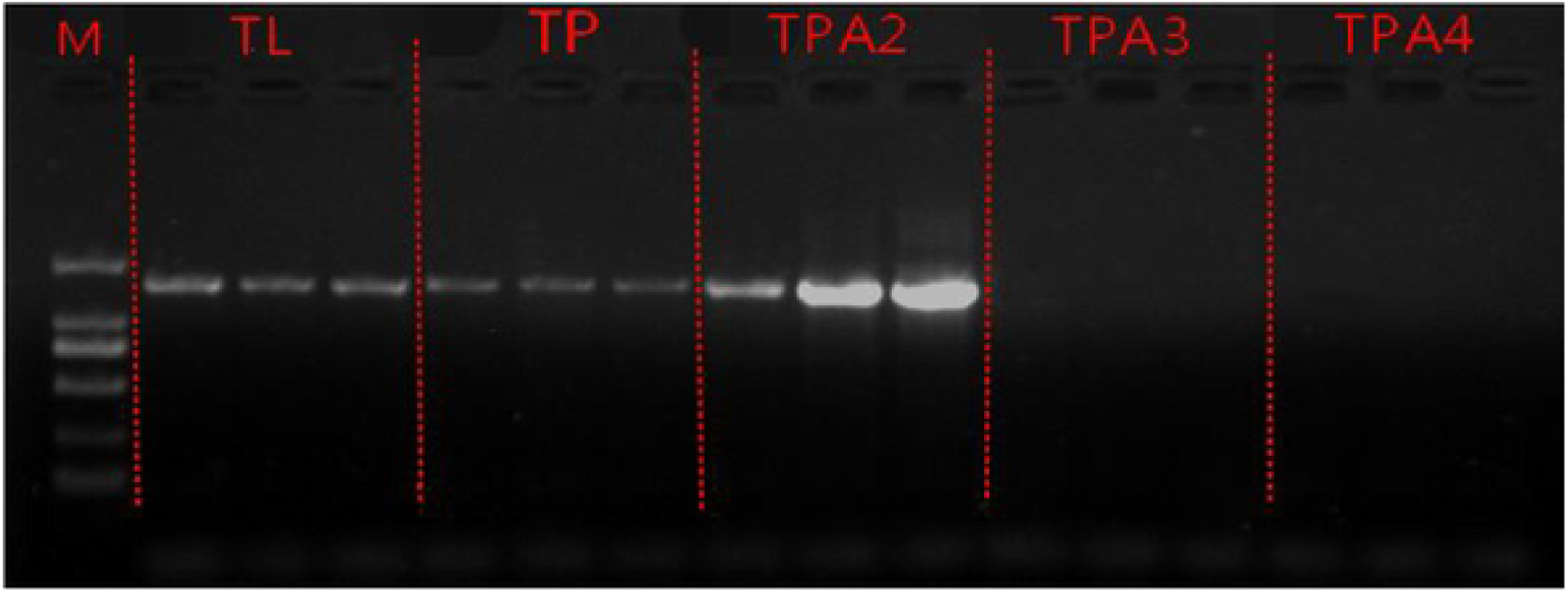
PCR Confirmation of reduced gut bacteria in *E.obliqua*. M, Marker. TL, Larvae reared normally on a tea leaf diet without antibiotics. TP, Larvae reared normally on a tea powder diet without antibiotics. TPA2, Larvae reared normally on a tea powder diet with antibiotics (200µ1/ml). TPA3, Larvae reared normally on a tea powder diet with antibiotics (300µ1/ml). TPA4, Larvae reared normally on a tea powder diet with antibiotics (400µ1/ml).

Culturing 4th-instar *E.obliqua* larvae gut homogenates on LB plates showed that treatment with antibiotics almost completely eliminated the cultivable bacterial in the gut of *E.obliqua* compared with the groups that received TL or TP only (Fig 3). Quantitative real-time PCR analysis showed that 16S rRNA gene expression in the TPA2 group was significantly higher (715.78 ± 274.78-fold) than in TPA3 (300μg/ml)antibiotics group (Student’s t-test, P < 0.001), indicating that almost no gut bacteria were present in the TPA-treated larvae. Next, we assessed the toxicity of the antibiotics to the insects.

**Figure 3.**
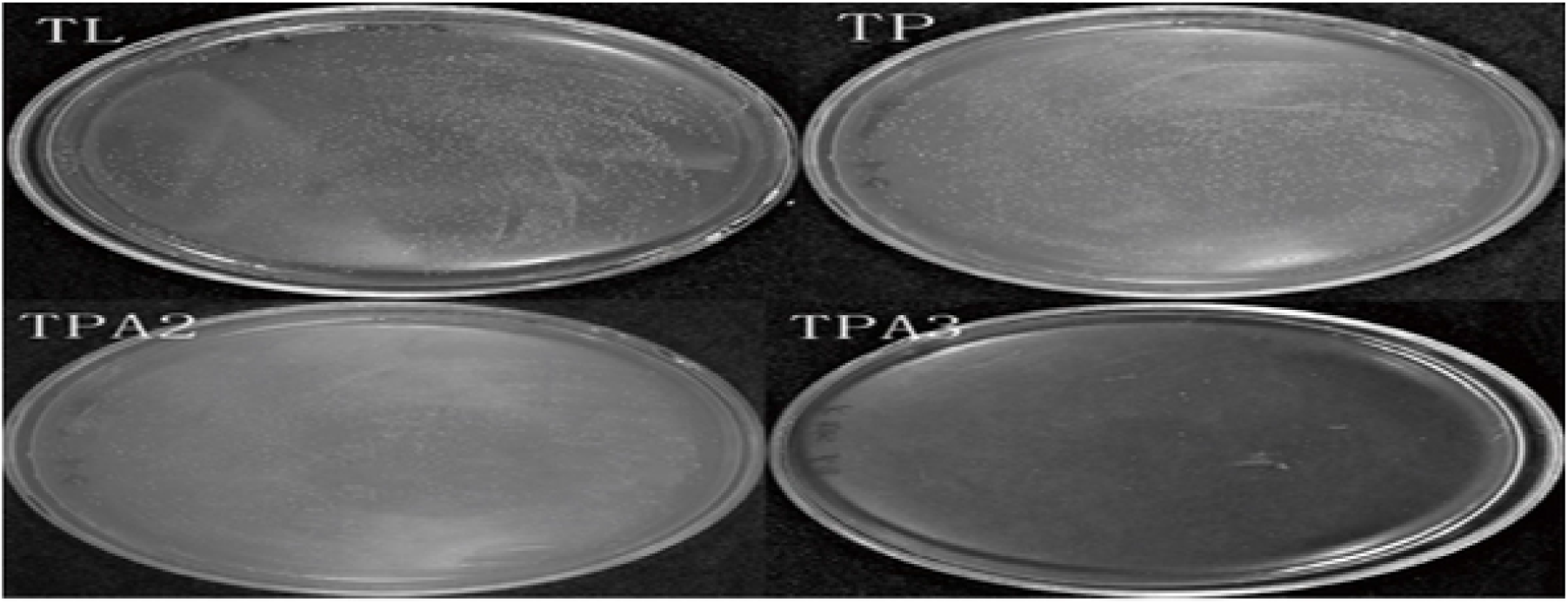
Effect of three different diets on culturablc gut bacteria in *E.obliqua*. TL, *E.obliqua* reared on a tea leaf diet. TP, *E.obliqua* reared on a tea powder diet. TPA2, *E.obliqua* reared on a tea powder diet containing antibiotics (200µg/ml). TPA3, *E.obliqua* reared on a tea powder diet containing antibiotics (300µg/ml).

### Effects of antibiotics on *E.obliqua* growth and reproduction

Comparing the four treatment diets had an effect on egg, larval, pupa and adult development times of *E.obliqua*. The results showed that treatment with 200 or 300 μg/ml of the antibiotic mixture had no significant effect on *E.obliqua* development time (egg, larvae, adult, pupa) compared with the antibiotic-free diet. However, comparing with the tea leaves group, the tea powder group and the antibiotic treatment groups exhibited significantly accelerated development, the larval and pupal of *E.obliqua* development times were obviously reduced (Fig 4). There was no significant difference between the weights of the female pupae and the male pupae in the four groups, but no matter which groups, the female pupae were heavier than the male pupae (Fig 5). Next, the eclosion rate, hatching rate, pupation rate, and fecundity rate were compared between the four groups. There was no significant difference in eclosion rate, egg hatching rate, or pupation rate between the groups that received antibiotics and those that did not, indicating that antibiotics had no significant effect on these indicators. The largest oviposition number of tea powder group was compared with other groups. However, the treatment with antibiotic mixture made the oviposition number notably reduced, but it did not effect the entire growth cycle (Fig 6). So antibiotic treatment was significant effected on fecundity of *E.obliqua*.

**Figure 4.**
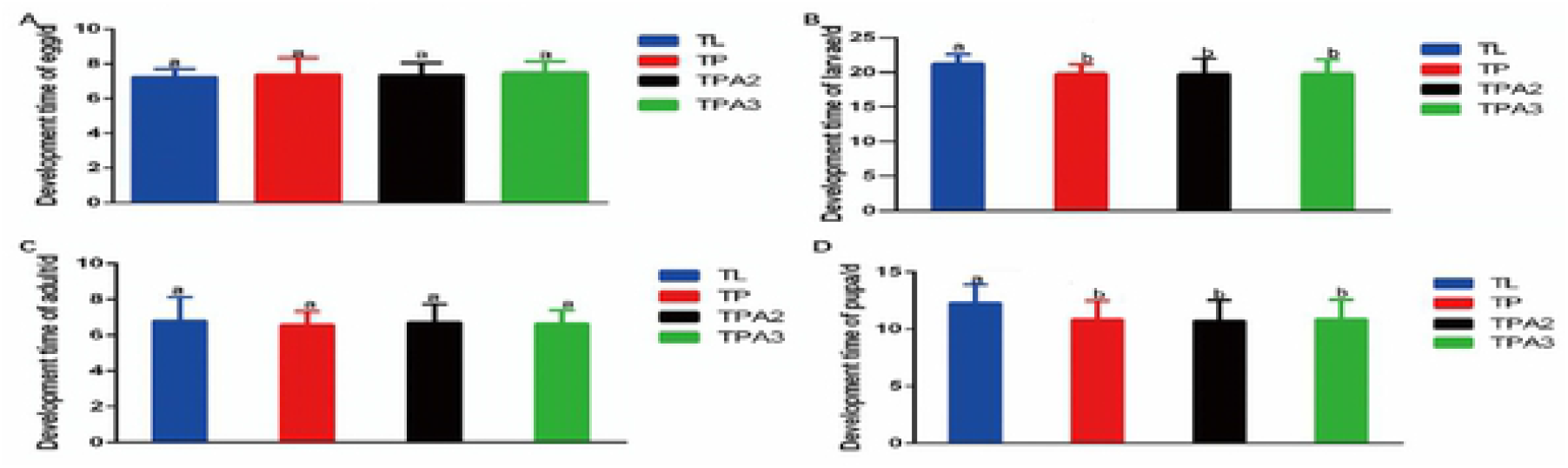
Impact of four different diets on *E.obliqua* development. (A) Impact of four different diets on *E. obliqua* egg development time.(B) Impact of four different diets on *E.obliqua* larval development time. (C) Impact of four different diets on *E.obliqua* adult development time. (D) Impact of four different diets on *E.obliqua* pupal development time.

**Figure 5.**
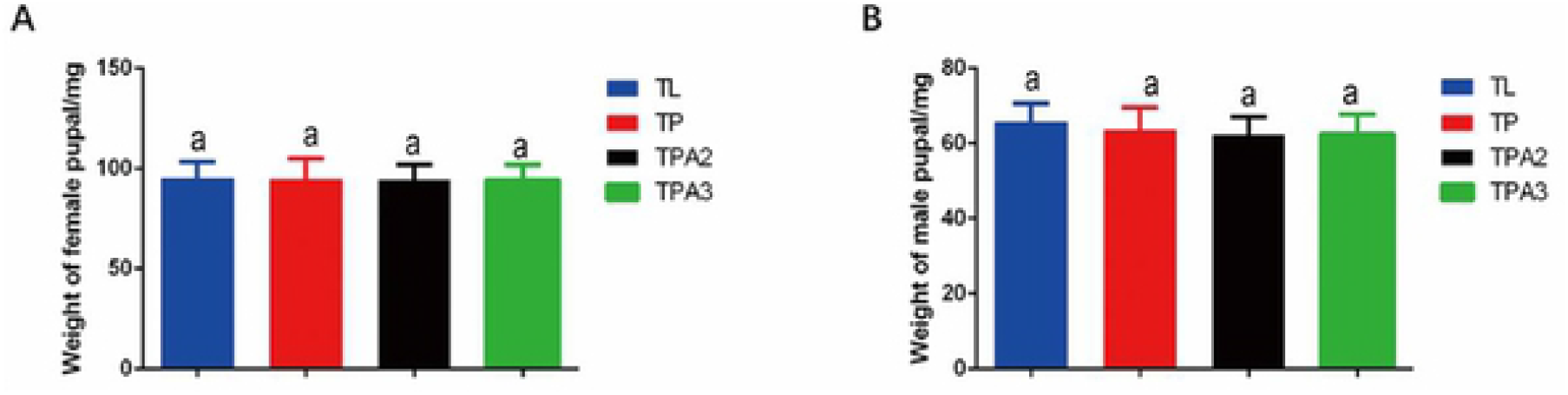
Impact of four different diets on *E.obliqua* pupal weight. (A) Impact of four different diets on *E.obliqua* female pupal weight. (B) Impact of four different diets on *E.obliqua* male pupal weight.

**Figure 6.**
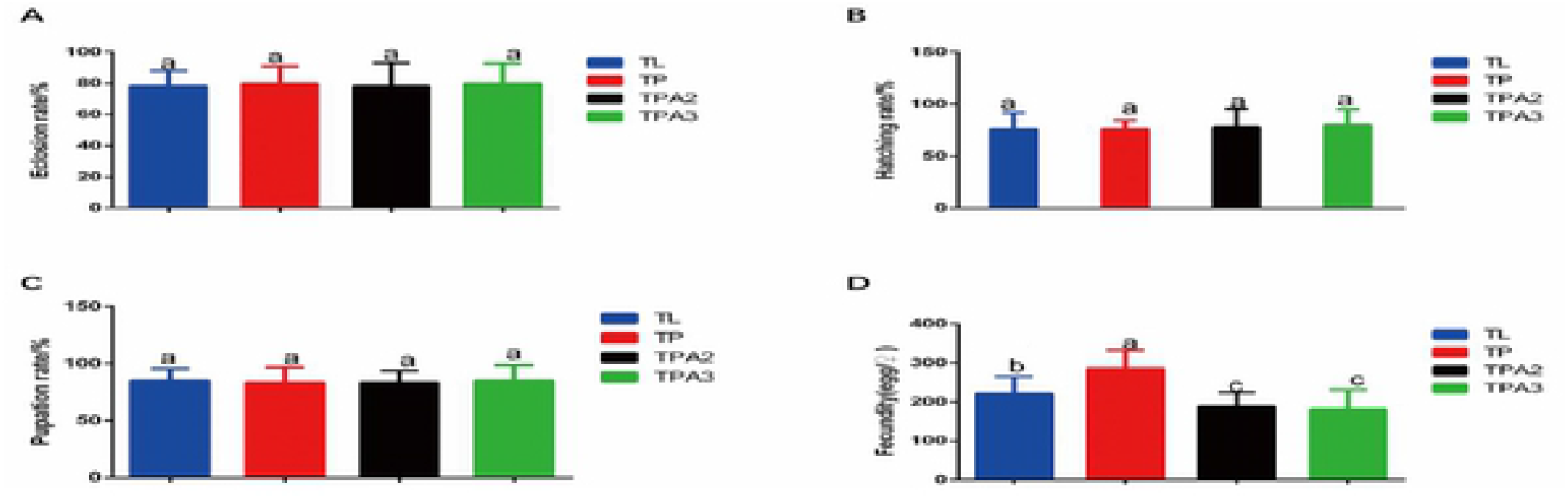
Impact of four different diets on *E.obliqua* reproduction. (A) Impact of four different diets on *E.obliqua* eclosion rate. (B) Impact of four different diets on *E.obliqua* egg hatching rate. (C) Impact of four different diets on *E.obliqua* pupation rate. (D) Impact of four different diets on *E.obliqua* fecundity.

## Discussion

Our results indicated that 300μg/ml of antibiotic mixture to treat *E.obliqua* could effectively remove gut microflora and obtain germ-free insect, as well as the treatment had no effect on growth and development of *E.obliqua*. Survival data were recorded for *E.obliqua* raised on diets containing different antibiotic concentrations. The mortality of larvae treated with 400μg/ml of the antibiotic mixture was high, while the other treatments had no effect on larval mortality. There was no significant difference in the growth and reproductive indicators that were assessed between the groups that received 200μg/ml or 300μg/ml of the antibiotic mixture, but there were significant differences in pupal and larval development times, as well as in fecundity. The larval and pupal development times in the group that received tea powder were significantly shorter than those in the tea leaf group. More fertile moths were found in the group that received tea powder compared with the tea leaves group, however, fewer fertile moths were found in the group that received antibiotics compared with the leaves and powder groups. The colony-forming units and PCR results showed that treatment with 200μg/ml of the antibiotic mixture does not effectively eliminate gut bacteria in *E.obliqua*, and may cause the gut bacterial community to become disturbed or even increase. Treatment with 300μg/ml of the antibiotic mixture effectively eliminated *E.obliqua* gut bacteria, so this concentration was selected for clearance of *E.obliqua* gut bacteria. Pupal and larval development time in tea powder group were significantly shorter than that in the tea leaves group. It has previously been reported that gut microbes are involved in cellulose degradation by insects [24]. However, a similar study of *P. xylostella*[12] concluded that a tea powder diet could be easily digested without the help of gut bacteria, so treatment with antibiotics did not affect normal feeding and larval development.

Sex-based differences in the insect responses to antibiotics is a very interesting research direction. When *Bemisia tabaci* (subtype Q) are treated with rifampicin, the incompatibility mainly exists in females. Studies had shown that *Wolbachia* affected host fecundity [25]. Our study confirmed that antibiotic-mediated gut bacteria clearance had a significant effect on *E.obliqua* fecundity, comparing with control groups without antibiotic. It may be that *Wolbachia* was eliminated by the antibiotic treatment, which significantly reduced the fecundity of *E.obliqua*. Previous studies that used antibiotics to eliminate gut bacteria[15, 16, 18] mainly focused on the effect of the antibiotics, but few of these studies investigated whether these antibiotics were toxic before they were applied. Our results showed that a specific concentration of the antibiotic mixture could be used to effectively eliminate gut bacteria with almost no effect on the larvae, so obtaining sterile insect.

## Acknowledgment

This study was financially supported by the National Natural Science Foundation of China (grant no.31870635 and 32072421), the Anhui Provinical Important Science & Technology Specific Projects (201903a06020019), Anhui Key Research and Development Program (202004e11020006) and National Key Research and Development Programs (2019yfd1001601).

